# Collection of continuous rotation MicroED Data from Ion Beam Milled Crystals of Any Size

**DOI:** 10.1101/425611

**Authors:** Michael W. Martynowycz, Wei Zhao, Johan Hattne, Grant J. Jensen, Tamir Gonen

## Abstract

Microcrystal electron diffraction (MicroED) allows for macromolecular structure solution from nanocrystals. To create crystals of suitable size for MicroED data collection, sample preparation typically involves sonication or pipetting a slurry of crystals from a crystallization drop. The resultant crystal fragments are fragile and the quality of the data that can be obtained from them is sensitive to subsequent sample preparation for cryoEM as interactions in the water-air interface can damage crystals during blotting. Here, we demonstrate the use of a focused ion beam to generate lamellae of macromolecular protein crystals for continuous rotation MicroED that are of ideal thickness, easy to locate, and require no blotting optimization. In this manner, crystals of nearly any size may be scooped and milled to ideal dimensions prior to data collection, thus streamlining the methodology for sample preparation for MicroED.

## Main Text

The cryoEM method microcrystal electron diffraction (MicroED) is a relatively new method with which structures of macromolecular assemblies are solved to atomic resolution from nanocrystals about one billion times smaller than what is typically used for X-ray crystallography^1–4^. These nanocrystals either grow spontaneously in sparse matrix crystallization trays or, when large crystals are obtained, can be sonicated or broken down to small crystallite fragments that are then used for MicroED^5^. Once good samples are available, grid preparation for cryoEM is arguably the most critical step before high-quality data can be collected. Typically, the crystallite solution is pipetted onto the holey carbon grid, excess solution is blotted away and the preparation plunged into ethane to create a layer of thin, amorphous ice embedding the crystallites. Many things can go wrong in the process. For example, the blotting might mechanically destroy the crystals; the blotting could be inefficient leading to very thick or very thin ice or lots of ice contamination on the grid; finally, the ice thickness can affect the hydration of the crystals and may deteriorate the underlying crystalline lattice^6,7^.

Some MicroED was collected recently from milled, fragmented crystals^8^. In that report, nano crystals were prepared as typical for MicroED^1,2,5^ after which the crystals were milled to thin lamellae and used for MicroED still diffraction experiments. Because only still diffraction^1^ was used the data had very low completeness and the resulting structure refined with poor statistics^8^. Importantly, the observation was made that the milling did not appear to severely damage the underlying lattice.

Here we demonstrate that large crystals, several hundred micrometers in thickness, can be scooped directly onto an EM grid, some excess solution gently blotted manually before flash freezing in ethane. The surrounding ice and embedded crystals are then milled using an ion beam down to any desirable thickness for analysis by MicroED. We show that by using this approach we could solve the structure of proteinase K from a single milled crystal. This approach considerably widens the scope of crystals suitable for MicroED, streamlines sample preparation, and decreases the time spent screening for crystals on the grid.

Large proteinase K crystals were grown by vapor diffusion as typically done for X-ray crystallography^9^. We took an entire 4μL drop of crystals and transferred all of it onto a freshly glow-discharged holey carbon grid. These grids were vitrified by plunging into supercooled ethane, and transferred to an FEI Versa FIB/SEM for milling without platinum coating. An overview of the grid in the SEM and by the FIB showed the grid overlaid with many large crystals of varying size (between ~5-300μm) (**Figure 1A-C**). An ideal specimen for milling was identified in low magnification SEM and FIB imaging by looking for a very large, sharp-edged crystals surrounded by amorphous ice that was located more than 10 μm away from the copper grid bar (**Figure 1B,C**). Crystals like this were easily found on every TEM grid tested, allowing us to mill several lamellae on each grid. It should be noted that for this demonstration we were deliberately targeting very large crystals that typically would not be amenable for analysis by MicroED (For example Figure 1C, arrow).

**Figure 1.**
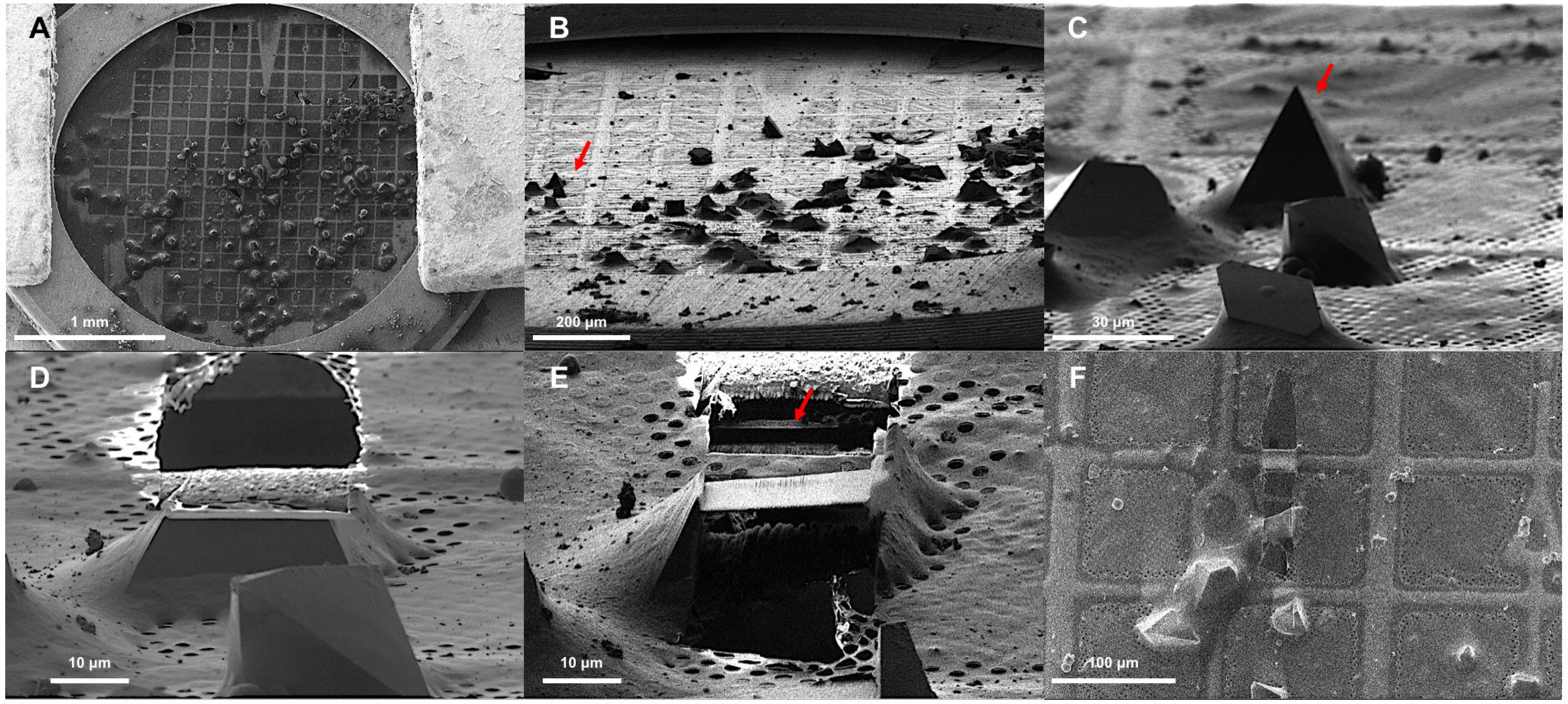
Preparation of macromolecular protein crystals for MicroED by focused ion beam milling. **A.** SEM image of a 3mm TEM grid. **B.** FIB image of the grid at 20^°^. **C.** FIB image of select crystals at high magnification. The arrow indicates the crystal that was milled. **D.** FIB image of select crystal from C after milling the top of the crystal lamella. **E.** FIB image of crystal after milling and cleaning both the top and bottom of the crystal leaving a lamella 300nm thick indicated by an arrow. **F.** SEM image of the crystal lamella at high magnification.

For milling we used a focused beam of gallium ions accelerated at 30kV. We initially gross-milled the surrounding media and the excess crystalline area using a beam current of 300 pA to clear away the large, thick, unwanted volumes above or below the sample to a thickness of approximately 1-3 μm (**Figure 1D,E**). Subsequent reductions in the size of the crystal lamella came by milling away equal volumes above and below the initially milled volume with reduced current in a step-by-step fashion as reported before^10,11^. The final lamella thickness was approximately 300nm with the final volumes removed with a gallium beam current of 30 pA (**Figure 1E**). The process of milling was monitored with SEM operated at 10 kV and 27 pA (**Figure1F**). The total time to mill a crystal of this size is approximately 10 minutes.

Following milling, the grids were transferred to an FEI Talos Artica 200kV electron microscope equipped with a bottom mount CetaD CMOS detector for MicroED data collection. Location of the crystalline lamellae after milling was apparent in low-magnification images by direct search or the inspection of a grid atlas (**Figure 2A**). The milled lamellae clearly stood out compared with un-milled regions and the milled crystals appear as a dark particle surrounded by the bright milled region (**Figure 2A**, **arrow**).

**Figure 2.**
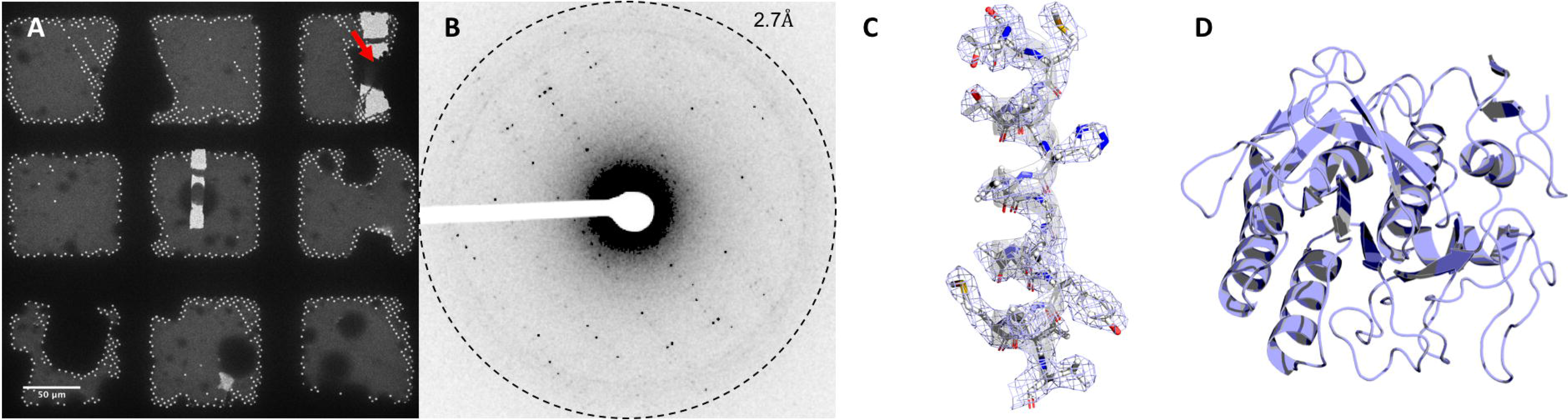
MicroED structure of proteinase K determined from a single milled crystal. **A.** Identification of milled crystals in low magnification TEM. Two lamella are easily identified, with an arrow indicating crystal from **Figure 1**. **B.** Single diffraction pattern from crystal in (**Figure 1/2A**). **C.** 2F_o_-F_c_ map of the central helix (residues 223-240) contoured at 1.5σ from the mean with a 2Å carve from the atomic centers for clarity. **D**. Final structure of proteinase K.

MicroED data were collected as previously described^2,12^. The milled crystals yielded diffraction to ~2.5Å resolution (**Figure 2B**) indicating that the milling process did not damage the underlying crystalline material. A complete data set was collected from a single milled crystal by continuous rotation MicroED, where a single lamella was continuously rotated while the diffraction was collected as a movie on the CetaD detector. The total exposure during the MicroED experiments was approximately 4 e^-^ A^-2^ (total dose) with a dose rate of 0.02e^-^ A^-2^ s^-1^ and a rotation speed of 0.3 degrees per second. The entire data set was collected in less than 5 minutes. The structure of proteinase K was solved by molecular replacement and refined to a final resolution of 2.75Å with acceptable R_work_ and R_free_ of 23% and 28%, respectively. A full description of data collection and statistics of the model is given in **Table 1**, and in the **Methods** sections.

**Table 1.** Data collection and analysis of a single, milled crystal

| Data | Parameter | Measure |
| --- | --- | --- |
| Data collection | Accelerating voltage (kV) | 200 |
|  | Electron source | Field emission gun |
|  | Total accumulated exposure (e<sup>-</sup>) | 4 |
|  | Crystals (#) | 1 |
|  | Microscope | Thermo-Fischer Talos Artica |
|  | Camera | Ceta-D |
|  | Rotation rate (°/s) | 0.297 |
|  | Wavelength (Å) | 0.0251 |
| Data analysis | Resolution range (Å) | 44.86 - 2.75 (2.848 - 2.75) |
|  | Space group | *P* 4<sub>3</sub> 2<sub>1</sub> 2 |
|  | Unit cell (a, b, c, α, β, γ) | 69.49 69.49 109.89 90 90 90 |
|  | Total reflections (#) | 31156 (3145) |
|  | Multiplicity | 4.7 (4.9) |
|  | Completeness (%) | 88.75 (89.61) |
| | Mean *I*/$\sigma$(*I*) | 2.93 (0.98) |
|  | Wilson B-factor | 36.4 |
|  | R-merge | 0.408 |
|  | R-pim | 0.2085 |
|  | CC1/2 | 0.92 |
|  | CC* | 0.979 |
|  | R-work | 0.2306 |
|  | R-free | 0.2780 |

The milling process did not appear to severely damage the underlying structure of the crystalline lattice. The only electron dose potentially absorbed by the lamella during FIB/SEM comes from the final image taken at low magnification used to view the lamella (**Figure 1F**). This dose should be surface-limited as the penetration depth of ~5keV electrons is only a few nanometers, suggesting that any irradiation by low energy electrons prior to milling is irrelevant^13–15^. Effects due to the focused ion beam imaging or the stress from physically milling away portions of the crystal did not manifest themselves in the final density. However, these effects may have led to a reduction of the observed resolution, or global damage to the lattice, limiting the resolution to ~2.7Å whereas previously we determined the structure of proteinase K by MicroED to 1.8Å resolution^5^. Gallium is expected to absorb into solid surfaces at grazing incidence angles, and might have absorbed into the crystal similarly to gas or liquid phase soaking in isomorphous replacement experiments^16^. Gallium was not observed in the final density map. However, this may be a possibility for thinner lamellae. Both the effects of radiation damage and absorption of gallium into the crystal should be investigated in the future.

In this work, we solved the structure of proteinase K by continuous rotation MicroED using a single crystal lamella milled by a gallium ion beam to a thickness of 300nm. Our results show that solving complete structures from single crystals milled by a FIB is possible and that the underlying crystalline material was not severely affected by the technique. This study broadens capabilities of MicroED because crystals that are too fragile to survive fragmentation methods or harsh blotting conditions could become amenable for data collection after FIB milling. Likewise, milling of crystals grown in lipidic cubic phase or other viscous solvents may now be possible. We believe this method greatly increases the potential scope of what can be done with MicroED, and adds a level of tunable control to all future MicroED experiments.

## Acknowledgements

The Gonen lab is supported by funds from the Howard Hughes Medical Institute. Work done in the Jensen lab was supported by NIH grant R35 GM122588. We would like to thank Ikaasa Suri for the help with sample preparation.

## Author Contributions

TG and GJ conceived the project. MWM prepared the protein crystals. WZ and MWM collected the FIB/SEM data and milled the crystals. MWM collected the MicroED data. MWM and JH processed the data. The manuscript was written by MWM and TG with contributions from all authors. MWM, GJ and TG designed the experiments.

## Protein Crystallization

Proteinase K (*E. album*) was purchased from Sigma and used without further purification. Crystals were grown in 24-well trays by sitting drop vapor diffusion using micro bridges (Hamilton). Drops were formed by combining 2μL of mother liqueur (1.25M Ammonium Sulfate 50mM Tris-HCl pH 8.5) and 2μL of protein solution (20 mg ml^-1^ Proteinase K from *E. album* 50mM Tris-HCl pH 8.5).

Protein crystals were taken directly from the drop using a 10μl pipette with the tip cut to increase its width. Approximately 4μl of protein solution was applied to the carbon side of a Quantifoil R 2/2 200 mesh finder grid (Ted Pella) in 100% humidity inside an FEI Vitrobot Mark IV after glow discharging for 60s at 15μA. Excess water on the grids was removed by gently pressing filter paper to the copper back side for 10s. Grids were then plunged directly into super cooled ethane without further blotting. Grids in ethane were transferred to liquid nitrogen and stored in a dewar until use.

## Scanning electron microscopy (SEM) and focused ion beam (FIB) milling

Grids were clipped and loaded into a FEI Versa FIB/SEM with a cryo-transfer system (PP3010T, Quorum Technologies). During FIB milling and transfer, samples were kept at liquid nitrogen temperature. SEM and FIB images were taken at various magnifications using dwell times between 100ns and 1μs. For milling, the gallium beam was accelerated by a voltage of 30kV and the stage was tilted at angles of 17°-20°. The milling current was gradually reduced from 300pA in the first round to 30pA as lamellae were thinned to their final thicknesses. SEM with 10 kV and 27 pA was used to monitor the milling progress.

## MicroED data collection and processing

Grids were loaded from the FIB/SEM to an FEI Talos Artica operating at liquid nitrogen temperatures. Lamellae were identified in low magnification imaging using the low-dose control at a typical magnification of 100× LM mode with a spot size 11. After identification of lamellae, continuous rotation MicroED data was collected as previously described^1,12,17^. Briefly, a selected area aperture was used to minimize background noise, and the stage was rotated at a constant rate of 0.30 degrees per second while a parallel beam of electrons scattered from the sample.

Data was converted from FEI’s SER format to SMV using in-house developed software available from https://cryoem.ucla.edu/downloads/snapshots. SMV images were indexed and integrated with XDS^18^; scaling and merging was performed using XSCALE^19^. The merged dataset was phased by molecular replacement in Phaser^20^ with 5i9s as the search model^9^, resulting in a final TFZ score >33 and LLG >1000. The structure was refined with phenix.refine using electron scattering factors to a final R_work_/R_free_ of 23/28% at a resolution of 2.75Å^21^.

## Statistics

Number values used to calculate the statistics in Table 1 were given by the total number of reflections and the number of unique reflections. No other statistical tests were used within the manuscript.

## Software

Figures were generated using PyMol (Schrodinger LLC) and ImageJ (NIH), and assembled in PowerPoint (Microsoft).

## Data Availability Statement

Atomic coordinates and structure factors were deposited to the Protein Data Bank (PDBID here) and the Electron Microscopy Data Bank (EMDB here).

